# A Phylogenetic Analysis of Shape Covariance Structure in the Anthropoid Skull

**DOI:** 10.1101/090910

**Authors:** Guilherme Garcia, Felipe Bandoni de Oliveira, Gabriel Marroig

## Abstract

Phenotypic traits evolve in a coordinated manner due to developmental and functional interactions, mediated by the dynamics of natural selection; the dependence between traits arising from these three factors is captured by genetic (**G**) and phenotypic (**P**) covariance matrices. Mammalian skull development produces an intricate pattern of tissue organization and mutual signaling that integrates this structure, although the set of functions it performs is quite disparate. Therefore, the interplay between these interactions, and their relationships with the adaptive landscape may thus influence divergence in covariance structure among sister lineages. Here, we evaluate the stability of phenotypic covariance structure in skull size and shape along the diversification of Anthropoid Primates under a explicit phylogenetic framework. We estimate diversity in covariance structure, testing hypotheses concerning the phylogenetic distribution of **P**-matrix variation and pinpoint which traits are associated with this variation. We find that most changes occurred in the basal split between Platyrrhini and Catarrhini, and that these changes occurred within both Orbital and Basicranial trait sets, while Oral, Nasal and Vault trait sets present stable associations along the Anthropoid phylogeny. Therefore, changes in **P**-matrix structure among Anthropoids are restricted to trait sets whose functional significance is associated with the accommodation of the two precursor tissues that compose the skull, while the stability in the remaining regions hints at the stability of the underlying functional relationships imposed by the adaptive landscape.

## Introduction

Phenotypic traits evolve in a coordinated manner either because of shared genetic and developmental processes or joint effects on fitness (Gould & Lewontin, 1979; Lande, 1979; Felsenstein, 1988; Zeng, 1988), and response to natural selection is maximized when both factors (variation and selection) are aligned, while discordance between them may deflect evolutionary response away from maximum increase in fitness (Schluter, 1996). The additive genetic covariances among traits (G) and the partial regression coefficients of fitness on traits (ft) provide linear approximations for both these effects in the characterization of phenotypic change across generations (Lande, 1979; Rice, 2002). Therefore, the additive covariances in G represent the codependency between traits due to pleiotropy and linkage disequilibrium, characterizing a linear genotype/phenotype map centered on the mean phenotype (Wagner, 1984,1996; Cheverud, 1996a).

However, the structure of pleiotropic interactions depends on the local curvature of the genotype/phenotype map, traditionally represented in quantitative genetics as either dominance or epistasis (Rice, 1998, 2004; Wolf *et al.*, 2001). The effect of epistatic *loci* on the modulation of pleiotropic interactions has been identified in experimental settings (e.g. Cheverud *et al.*, 2004; Wolf *et al.*, 2005; Pavlicev *et al.*, 2008), indicating that populations may harbor genetic variation in the association between traits. Genetic covariances among phenotypic traits then evolve as a consequence of changes in allele frequencies in these *loci*, for example in response to genetic drift and founder effects (Goodnight, 2000; Brito *et al.*, 2005). Local features of the adaptive landscape may also impact genetic covariances among traits, as G is thought to match the patterns imposed by stabilizing selection and mutational effects (Lande, 1980; Cheverud, 1984; Jones, 2007). It is noteworthy that changes in genetic covariances due to the curvature of adaptive landscapes can be explained just by considering shifts in linkage disequilibrium among *loci* (Turelli, 1988), without need to appeal to epistatic pleiotropy. However, recent experimental data (Careau *et al.*, 2015) and simulation-based models (Jones *et al.*, 2012; Melo & Marroig, 2015) have demonstrated the effects of directional selection on the structure described by G, thus indicating that the linear component of adaptive landscape can also have an impact on genetic covariances due to genetic variation in pleiotropic interactions.

The relationship between adaptive landscapes and intrapopulational covariance structure is mediated through performance (Arnold, 1983), which may be thought as a dynamical property of phenotypes. This implies that the separation between developmental and functional interactions as two distinct factors shaping phenotypic covariance structure is blurry at best (Cheverud, 1996a; Zelditch & Swiderski, 2011). For example, in the mammalian skull, precursor tissues originate two distinct regions, Face and Neurocranium, which exhibit marked contrasts in terms of functions which they perform, interactions with soft tissues, and response to developmental milestones, such as birth or weaning (Hallgrímsson & Lieberman, 2008; Lieberman, 2011). Thus, the Oral, Nasal, and Zygomatic regions are associated with the Face and are responsible for mastication, respiration, and the attachment of the muscular apparatus involved in mandibular articulation. The Vault, Orbit and Base regions are associated with the Neurocranium and are responsible for encasing and protecting both brain and eye, and for supporting and connecting the skull with the rest of the body. The Vault also houses muscle attachment sites associated with mandibular articulation, indicating that, to some degree, regions may be involved in more than one function and that some functions may be shared between them. The contrast between these regions is the result of distinct gene expression profiles, which are further changed by the diffusion of signaling factors, thus generating a feedback loop of cell and tissue differentiation (Turing, 1952; Marcucio *et al*, 2005; Meinhardt, 2008; Franz-Odendaal, 2011; Xu *et al*, 2015).

These signaling factors may target specific cell lineages, but the contact between neighboring tissues may produce correlated changes between them due to mechanical interactions and through mutual signaling cascades that induce specific behaviors, and such interactions are necessary for the proper development of both tissues (Cheverud *et al.*, 1992; Ravosa *et al.*, 2000; Jiang *et al.*, 2002; Marcucio *et al.*, 2011). For instance, cranial Vault growth, which occurs on the final stages of pre-natal development, is a result of the tension exerted by the growing brain on its encircling membranes, inducing them to secrete signaling factors which promote bone growth (Opperman, 2000; Rice *et al.*, 2003). This mechanism promotes a tight association between the Vault and the brain. Furthermore, post-natal facial growth is induced by muscular activity related to masticatory function, and this effect dominates post-natal skull growth (Zelditch *et al.*, 1992; Herring, 2011). Muscle activity mainly affects the focal Oral region in which muscular forces are exerted, but other skull regions to which these muscles are attached are also affected to a lesser extent, such as the Zygomatic and Vault. Therefore, development is composed of a series of such events, and it may be difficult to isolate the effect each individual process has on covariance structure, given their spatial overlap (Hallgrímsson *et al.*, 2009); hence, both the temporal hierarchy and the spatial organization of developmental and functional interactions influence the patterns embedded in genetic covariance structure.

Empirical evidence on long-term changes on genetic covariance structure (comparative quantitative genetics; Steppan *et al.*, 2002) rely on the correspondence between G and its phenotypic counterpart P, because estimating G demands high sample sizes and nonetheless such estimate is prone to substantial error, since sample units are families rather than individuals (Meyer, 1991; Houle & Meyer, 2015). The similarity in covariance structure between P and G (“Cheverud’s Conjecture”; Cheverud, 1988; Roff, 1995), has been supported in different trait systems and organisms (e.g.: Waitt & Levin, 1998; Dochtermann, 2011; Garcia *et al.*, 2014), indicating that environmental effects (E) are either uncorrelated or exhibit patterns similar to G, since these effects exert their influence upon phenotypes through the same developmental processes by which genetic variation is structured (Rice, 2002, 2004).

The comparative analysis of phenotypic covariance structure shows that P can remain stable in macroevolutionary scales (e.g. Marroig & Cheverud, 2001; Oliveira *et al.*, 2009b; Kolbe *et al.*, 2011). This stability may be a consequence of the alignment between phenotypic covariance structure and the local features of the adaptive landscape acting over different lineages. In New World Monkeys (Platyrrhini), divergence in body size among lineages is closely associated with shifts in diet composition (Rosenberger, 1992). Size represents the main feature of both genetic (Cheverud, 1995, 1996b) and phenotypic (Marroig & Cheverud, 2005) covariance structure, and for some groups within New World Monkeys, such as Atelids and Callithrichines, divergence in body size is a direct consequence of directional selection (Marroig & Cheverud, 2010; Marroig *et al.*, 2012). On the other hand, dietary or locomotory divergence between sister lineages can rearrange the patterns expressed in P (Young & Hallgrímsson, 2005; Young *et al.*, 2010; Monteiro & Nogueira, 2010; Haber, 2015), and such reorganization hints at changes in the underlying architecture of the genotype/phenotype map, indicating that the patterns privileged by selection can overcome constraints imposed by development (Jamniczky & Hallgrímsson, 2009).

The adaptive radiation of phylostomid bats, for instance, has involved rearrangements of mandibular phenotypic covariance structure (Monteiro & Nogueira, 2010), as the dietary divergence in this group from a probable insectivore ancestor towards more specialized diets (such as frugivory or sanguivory) imply distinct functional relationships among mandibular components. This reorganization may indicate heterogeneity in the non-linear aspects of the adaptive landscape for different phylostomid lineages. For bats in general, their specialized locomotory behavior is associated with the decoupling between fore- and hindlimb covariance structure when compared to other mammalian lineages (Young & Hallgrímsson, 2005). To a lesser extent, a similar pattern can be observed in hominoids when compared to remaining Anthropoids (Young *et al.*, 2010). Another example of divergence in covariance structure among lineages due to changes in functional demands are those observed in ruminants (Haber, 2015), which may related to increased metabolic requirements for occupying open habitats, as the divergence between Bovidae and Cervidae is associated with increased Nasal integration in the former. Thus, divergence among sister lineages may imply a reorganization of both genetic and phenotypic covariance structure of phenotypic traits if such divergence is associated with distinct functional relationships, represented by the non-linear components of adaptive landscape.

Currently, there are several methods dedicated to the comparison of covariance structures (e.g. Krzanowsky, 1979; Phillips & Arnold, 1999; Cheverud & Marroig, 2007), and such methods are often focused on constructing hypothesis of similarity or dissimilarity; However, it is not always clear which hypothesis is the adequate one, and frequently either hypotheses can be rejected for the same pair of matrices, hindering interpretation (Ackermann & Cheverud, 2000; Marroig & Cheverud, 2001; Haber, 2015). By expressing covariance matrix (dis)similarities with a single metric, these methods lack an explicit way of describing structural differences between pairs of covariance matrices. Furthermore, these methods also lack a direct manner to incorporate phylogenetic relatedness in pairwise comparisons. While extensions for these methods have been proposed to deal with the first limitation (e.g. Hansen & Houle, 2008; Hine *et al.*, 2009; Marroig *et al.*, 2011), the second issue is usually resolved by comparing the set of pairwise matrix comparisons with the set of phylogenetic distances among lineages (e.g. Marroig & Cheverud, 2001; Oliveira *et al.*, 2009b). Although such comparison provides a first approximation to this problem, it has the same problems as comparing covariance matrices themselves, that is, summarising the relationship between patterns expressed in P or G in a set of lineages and their phylogenetic relatedness to a single value. In this manuscript, we explore novel methods to circumvent these issues.

### Objectives

The interplay between developmental and functional interactions, and their relationships with the topology of adaptive landscapes may influence the divergence between covariance structure in sister lineages. In the present work, we evaluate the stability of phenotypic covariance structure in skull size and shape along the diversification of Anthropoid Primates. We build upon approaches proposed by other authors (Marroig *et al.*, 2011; Aguirre *et al.*, 2013; Haber, 2015) in order to explicitly incorporate phylogenetic relationships into the comparative analysis of covariance structure, under the hypothesis that different cranial regions will exhibit different degrees of stability among sister lineages, thus producing a non-random pattern of changes in covariance structure.

## Methods

### Sample

Our sample consists of 5108 individuals in 109 species, distributed throughout all major Anthropoid clades above the genus level, comprising all Platyrrhini genera and a substantial portion of Catarrhini genera. We associate this database with a ultrametric phylogenetic hypothesis for Anthropoidea (Figure S2), derived from Springer *et al.* (2012).

Individuals in our sample are represented by 36 landmarks; these landmarks were registered using a Polhemus 3Draw and a Microscribe 3DX for Platyrrhini and Catarrhini, respectively. Twenty-two unique landmarks represent each individual (Figure S1), since fourteen of the 36 registered landmarks are bilaterally symmetrical. For more details on landmark registration, see Marroig & Cheverud (2001) and Oliveira *et al.* (2009b). Databases from both previous studies were merged into a single database, retaining only those individuals in which all landmarks from both sides were present. In the present work, we considered only covariance structure for the symmetrical component of variation; therefore, prior to any analysis, we controlled the effects of variation in assymmetry. We followed the procedure outlined in Klingenberg *et al.* (2002) for bilateral structures by obtaining for each individual a symmetrical landmark configuration, averaging each actual shape with its reflection along the sagittal plane.

We used this database to obtain local shape variables (Márquez *et al.*, 2012), which represent infinitesimal volumetric expansions or retractions, calculated as the natural logarithm determinants of derivatives of the thin-plate spline between each individual in our sample and a reference shape (in our case, the mean shape for the entire sample, estimated from a Generalized Procrustes algorithm). Such derivatives were evaluated at the midpoints between pairs of landmarks represented in Figure S1, for a total of 38 local shape variables.

After obtaining these values, we estimated covariance P-matrices for size (represented by the natural logarithm of Centroid Size) and local shape variables after removing fixed effects of little interest here, such as sexual dimorphism, for example. These effects were removed through a multivariate linear model adjusted for each species, according to Figure S2. We adjusted such models under a Bayesian framework, sampling 100 residual covariance matrices from the posterior distribution of each model. These distributions allow us to estimate uncertainty for any parameters derived from these matrices as credibility intervals; furthermore, since posterior distributions are conditional upon the prior distribution we used – a uniform Wishart distribution – every matrix sampled from these posterior distributions is also a realization of a Wishart distribution, therefore positive-definite regardless of sample size (Gelman *et al.*, 2004). In this framework, lower sample sizes imply in broader and less informative credibility intervals. For each posterior sample, we estimated geometric mean covariance matrices, since this mean respects the underlying geometry of the Riemannian manifold in which positive-definite symmetric matrices lie (Moakher, 2005, 2006). These mean P-matrices are also positive-definite, regardless of sample size. For each species, we ran independent models, with 13000 iterations of MCMC sampling, discarding the 3000 initial iterations as a burn-in period and further sampling one covariance matrix per 100 iterations to avoid autocorrelations induced by sequential sampling.

## Phylogenetic Decompostion of Matrix Diversity

In order to evaluate the distribution of covariance structure diversity during Anthropoid diversification, we estimated Riemannian distances among all pairs of mean P-matrices, according to the definition given by Mitteroecker & Bookstein (2009); for any pair of positive-definite covariance matrices C_*i*_ and C_*j*_ of size *p* × *p*, the distance d(C_*i*_, C_*j*_) is given by

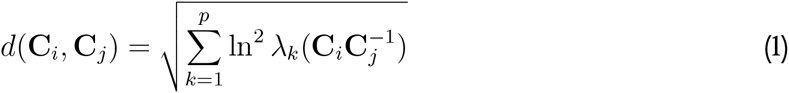

where *λ_k_(·)* refers to the *k*-th eigenvalue obtained from the spectral decomposition of a given matrix, in this case the product 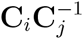. This distance among pairs of P-matrices is negatively correlated with Random Skewers comparisons (Figure S3), a measurement of matrix similarity explored elsewhere (Cheverud & Marroig, 2007). The similarity between Riemannian distances and Random Skewers similarity indicate that our conclusions would be the same regardless of the metric used to characterize matrix similarity or dissimilarity.

Using these distances among P-matrices, we estimate matrix diversity at each node of the phylogenetic tree of Anthropoidea using a measurement of the weighted distance among the distributions of matrix distances for the two descending edges, based on Pavoine *et al.* (2010). For a fully resolved tree, diversity *w_i_* on node *i* is given by

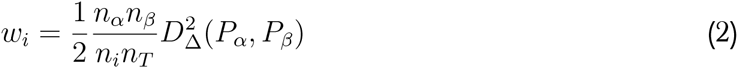

where *α* and *β* represent the subsets of descendants from node *i*, and *n* refers to the number of species on each set (*n_i_* for the total descendants of node *i*; *n_T_* for the total number of species considered; *n_α_* and *n_β_* for the size of descending subsets). *D_Δ_*(*P_α_*, *P_β_*) represents the actual distance between the two distributions *P_α_* and *P_β_* for descending nodes, as formulated by Rao (1982):

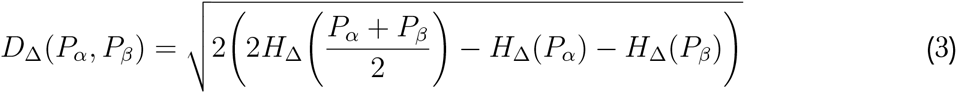

where

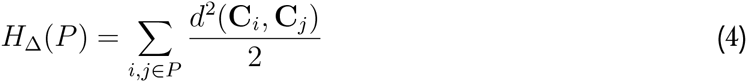

represents Rao’s quadratic entropy among Riemannian distances *d*(C_*i*_, C_*j*_) as defined in Equation 1.

Following the framework estabilished by Pavoine *et al.* (2010), diversity *w_i_* can be normalized as *v_i_ = w_i_/∑_i_ w_i_* to represent the percentage of diversity with respect to the total diversity on the phylogenetic tree. We test three different hypothesis regarding the distribution of *v_i_* values through Anthropoid diversification: (1) that P-matrix diversity is concentrated in a single node; (2) that P-matrix diversity is concentrated in a reduced number of nodes; (3) that P-matrix diversity is skewed towards either the root or tips of the phylogeny, in a two-tailed test. We test each hypothesis against the null hypothesis that the distribution of matrix diversity is randomly arranged over the phylogeny; such null hypothesis is represented by randomizing the association between terminal branches and covariance matrices, constructing 9999 distributions of *v_i_* values that represent this scenario. Each test is carried out using a different parameter derived from the distribution of *v_i_* values (described in detail by Pavoine *et al.*, 2010), comparing the actual value obtained with a null distribution constructed using permutations. The third hypothesis can be tested either by considering only the topology of the tree and by also considering branch lengths; both tests are similar to Blomberg’s (2003) *K* test, as they search for a phylogenetic signal in covariance structure diversity.

## Characterizing Covariance Matrix Variation

The tests described in the previous section allow us to pinpoint which nodes contribute mostly to divergence in covariance structure; however, these tests are not designed to properly describe the actual changes in P-matrix structure that are responsible for such divergence. To actually represent these changes in a comprehensible manner, we combine a number of ordination techniques to reduce the dimensionality of the manifold that contains covariance matrices of size *p × p* (Figure 1).

For a Riemannian manifold, there exists at least one bijective function defined in the neighbourhood of a given covariance matrix M that maps the manifold to an Euclidean space – a hyperplane with *p*(*p* − 1)/2 dimensions also contained in R^*p*×*p*^ – and equips the manifold with a notion of inner product, thus allowing the construction of an orthonormal basis that can be used to describe variation in P-matrix structure. For a covariance matrix X in the neighbourhood of M,

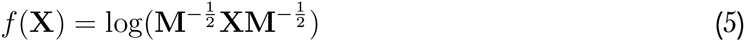

represents one possible function. Here, the logarithm operator refers to matrix logarithm; for symmetric positive-definite matrices, this transformation is equivalent to applying the usual logarithm function to the eigenvalues of such matrix and reverting the spectral decomposition. The function defined in Equation 5 also transforms the Riemannian distance among covariance matrices defined in Equation 1 into Euclidean distances between transformed matrices (Moakher, 2005).

We defined the average matrix among all sampled P-matrices as the location parameter M to map the entire set of posterior P-matrices into an Euclidean space. We then used these P-matrices to produce axes of matrix variation using an eigentensor decomposition (Basser & Pajevic, 2007; Hine *et al.*, 2009) obtaining a set of eigentensors and eigenvalues that summarise matrix variation (Figure 1a). As a consequence of using the mean covariance matrix for the entire sample over Equation 5, the projections over eigentensors we obtained are naturally centered on M.

We used the projections of P-matrices over these eigentensors as traits in a phylogenetic Principal Component Analysis (pPCA; Jombart *et al.*, 2010), which produces a new set of axes of matrix variation that considers both trait dispersal and phylogenetic relationships among species simultaneously. If Z represents a matrix with projections of *n* P-matrices over each eigentensor on its columns, phylogenetic PCs are the eigenvectors obtained from

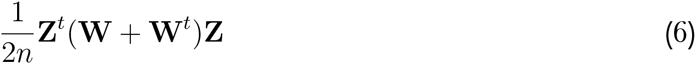

where W represents the matrix of phylogenetic distances between species; here, the distance *W_ij_* between tips *i* and *j* is the sum of branch lengths from their last common ancestor to both tips. Other definition of phylogenetic distances may be used (see Jombart *et al.*, 2010); the results we show here were not changed by considering different measures of phylogenetic distance, regardless of whether these distances consider branch lengths among species or not.

**Figure 1:**
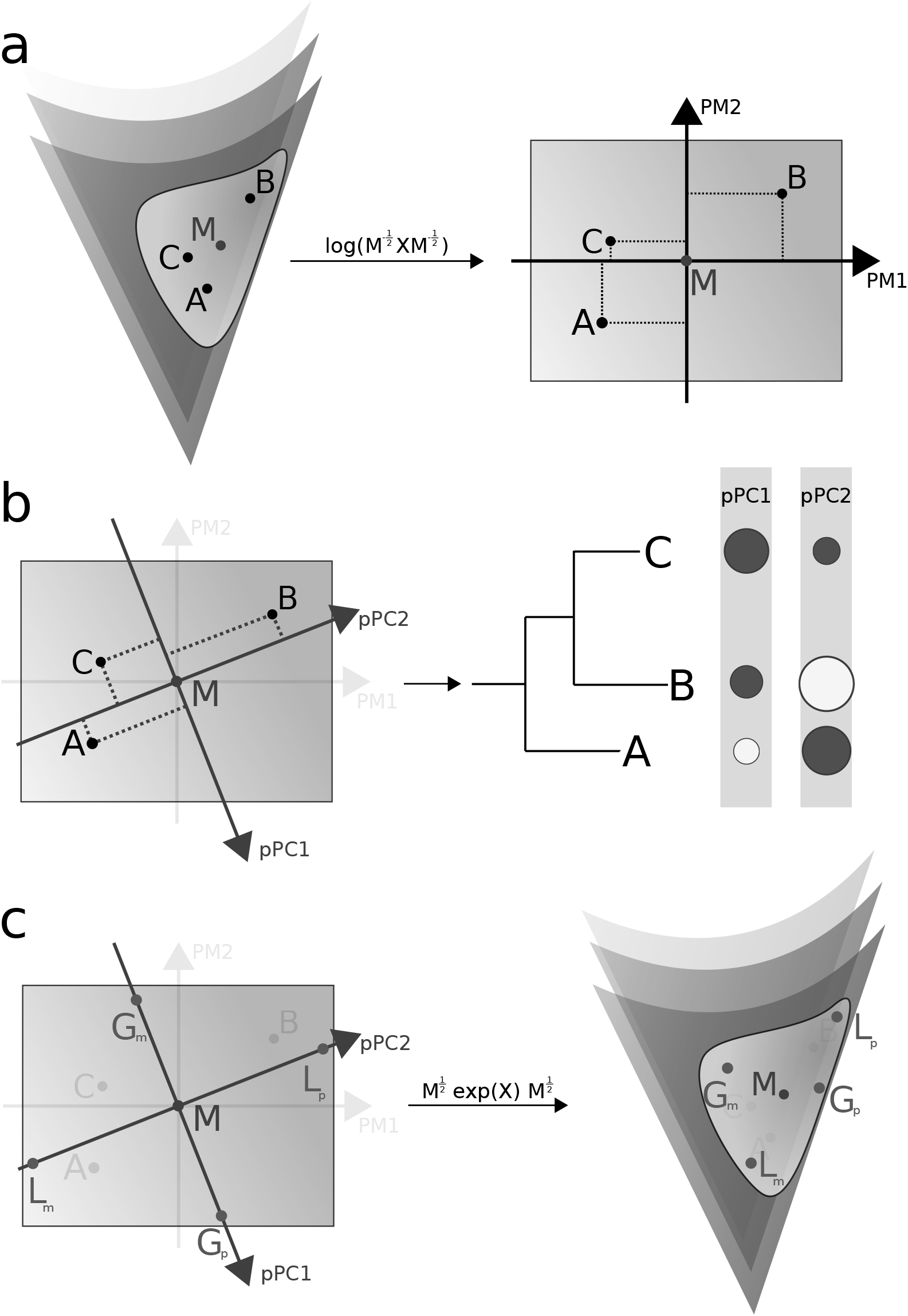
Representation of the steps used to characterize covariance matrix variation. In (a), the set of covariance matrices A, B and C in the neighbourhood of M are projected into an Euclidean space and eigentensors are estimated (PM1 and PM2); in (b), these eigentensors are rotated to incorporate phylogenetic relatedness; in (c), covariance matrices at the upper and lower bounds of the confidence intervals for each axis are returned back to the original manifold. See text for more details.

Such analysis produces both positive and negative eigenvalues, which are respectively associated with variation close to the root of the tree (‘Global’) and variation close to the tips (‘Local’) in matrix structure (pPC1 and pPC2 in Figure 1b, respectively). Pavoine *et al.* (2010) argues that this contrast between Global and Local components in phylogenetic PCs reflects phylogenetic signal and convergence in trait values, respectively, as observed in the distribution of Moran’s (1948, 1950) autocorrelation Indexes for each axis constructed in this manner. This index can be understood as the degree onto which an observed value in a given species is determined by the values on its phylogenetic “neighborhood” (as expressed by W), in a similar manner to autoregressive models (Cheverud & Dow, 1985; Cheverud *et al.*, 1985).

For each pPC obtained in this manner, we obtained two covariance matrices by estimating the upper and lower limits of the 95% confidence interval for each axis and mapping these values back to the manifold of symmetric positive-definite matrices (Figure 1c), defining the inverse operation associated with Equation 5 as

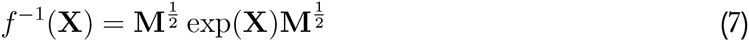

where the exponential operator refers to matrix exponential. We used these covariance matrices to describe matrix variation associated with each axis comparing each pair of matrices with the Selection Response Decomposition tool (Marroig *et al.*, 2011), in order to pinpoint which traits are associated with the divergence in covariance structure associated with each pPC.

In order to characterize such divergence in covariance structure with respect to the uncertainty in P-matrix estimation, we carried out the analyses described in this section with both mean P-matrices obtained from posterior samples, and with posterior samples themselves, obtaining 100 sets of phylogenetic PCs and 100 sets of paired covariance matrices for each axis, thus allowing us to estimate a posterior distribution of mean SRD scores for each trait in all pPCs. We use the phylogenetic PCA estimated over mean P-matrices in order to represent the phylogenetic patterns described by each pPC.

We use the posterior distribution of mean SRD scores over traits and pPC to investigate whether these changes in trait-specific covariance structure along Anthropoid diversification are randomly distributed with respect to the skull regions delimited in Table S2 by comparing SRD scores estimated within each region for all pPCs. The association between cranial traits and such regions reflect their functional significance and developmental origins.

### Software

We performed all analyses under R 3.2.1 (R Core Team, 2015). We fitted Bayesian linear models for estimating posterior P-matrix samples using the *MCMCglmm* package (Hadfield, 2010). Both eigentensor decomposition and the SRD method are provided by the *evolqg* package (Melo *et al.*, 2015); the phylogenetic decomposition of diversity was provided by Pavoine *et al.* (2010) in their Supplemental Material, while pPCA is implemented in the *adephylo* package (Jombart & Dray, 2010). In order to obtain symmetrical landmarks configurations, we used code provided by Annat Haber, available at http://life.bio.sunysb.edu/morph/soft-R.html.

## Results

The distribution of covariance matrix diversity along the Anthropoid phylogeny (Figure 2a) indicates that the divergence between Catarrhini and Platyrrhini contributes to approximately 10% of all covariance matrix diversity; within New World Monkeys, the divergence between Atelidae and Cebidae contributes with 4% to covariance structure diversity, while for Catarrhines, the divergence between Hominoidea and Cercopithecoidea contributes 3% to overall covariance matrix diversity. The remaining P-matrix diversity is distributed along the tree, with a consistent decay of explained diversity the closer any given node is from terminal branches. With respect to the hierarchy of tests exploring the phylogenetic distribution of matrix diversity (Table 1), all tests reject their null hypotheses of random arrangements of diversity along the Anthropoid phylogeny, thus indicating that covariance matrix diversity exhibits some degree of phylogenetic structure, with these few basal nodes contributing to a greater extent to such diversity.

**Table 1:** Phylogenetic decomposition of covariance matrix structure diversity.

|  | **Value** | **Expected<sup>a</sup>** | **Distance<sup>b</sup>** | ***p*-value** |
| --- | --- | --- | --- | --- |
| Single Node | 0.106 | 0.029 | 13.455 | $< 10^{-4}$ |
| Few Nodes | 0.248 | 0.139 | 13.545 | $< 10^{-4}$ |
| Root/Tip Skewness<sup>c</sup> | 0.632 | 0.505 | 12.197 | $< 10^{-4}$ |
| Root/Tip Skewness<sup>d</sup> | 0.381 | 0.505 | -11.067 | $< 10^{-4}$ |

<sup>a</sup> refers to the distribution of permuted values;
<sup>b</sup> difference between empirical and expected values in standard deviations of the permuted values distribution;
<sup>c</sup> considering only topology;
<sup>d</sup> including branch lengths.

The eigenvalue distribution of phylogenetic Principal Components that describe P-matrix structure (Figure 3) shows that individual Global components surpass their Local counterparts in terms of explained variance, such that the first Global component has a larger contribution than the first Local component to interspecific P-matrix variation. While there are less positive than negative eigenvalues, Global components explain more than half (57%) of the total P-matrix variation, considering mean absolute eigenvalues obtained from their posterior distribution. The posterior distribution of Moran’s Index for phylogenetic Principal Components (Figure 4) is assymetric towards positive values, also indicating that the similarity produced by phylogenetic inertia is greater than the similarity produced by convergence in P-matrix structure.

Considering the distribution of matrix projections for each species on these pPCs (Figure 2b), we observe that the first Global pPC separates New World and Old Monkeys, while the second Global pPC consists of a contrast between Atelids and Cebids, and the third Global pPC generally contrasts Hominids with the remaining Anthropoids, thus indicating a pattern of P-matrix variation consistent with those observed in the diversity decomposition summarized in Figure 2a. The first Local pPC consists of localized contrasts between sister species, such as the two representatives of *Saimiri*, for instance; however, matrix projections over this axis can be explained by the effect of sample sizes (Figure S4). These localized contrasts can thus be explained on the account of substantial sample size differences between sister species.

**Figure 2:**
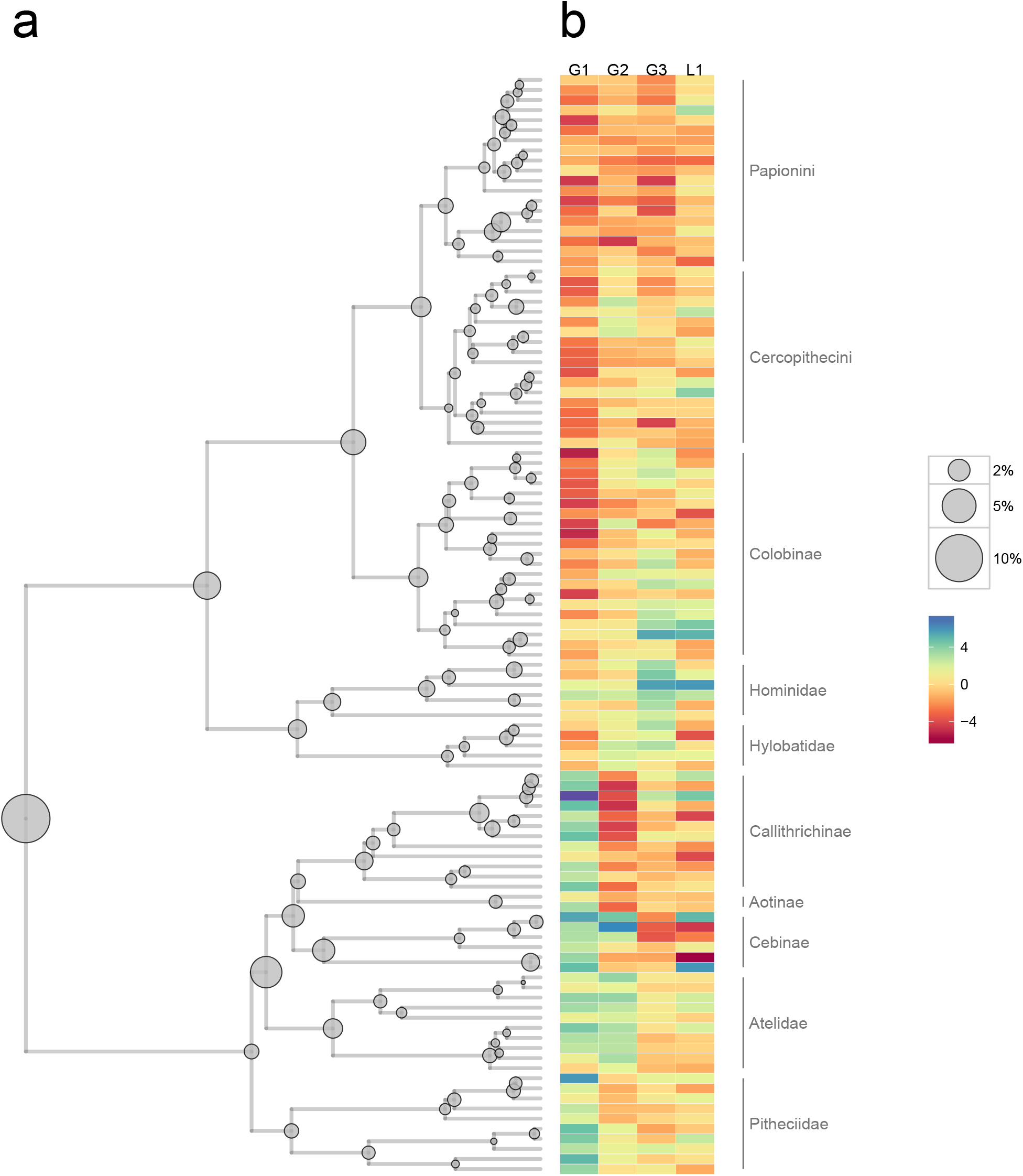
Phylogenetic decomposition of P-matrix variation. (a) Decomposition of matrix diversity over the phylogenetic hypothesis for Anthropoidea; the size of each circle indicates the percentage of diversity on each node, according to the legend. (b) Mean P-matrices of each species projected over the first three and the last phylogenetic Principal Components (G1-3 and L1, respectively); cell colors represent projection values, according to the legend.

**Figure 3:**
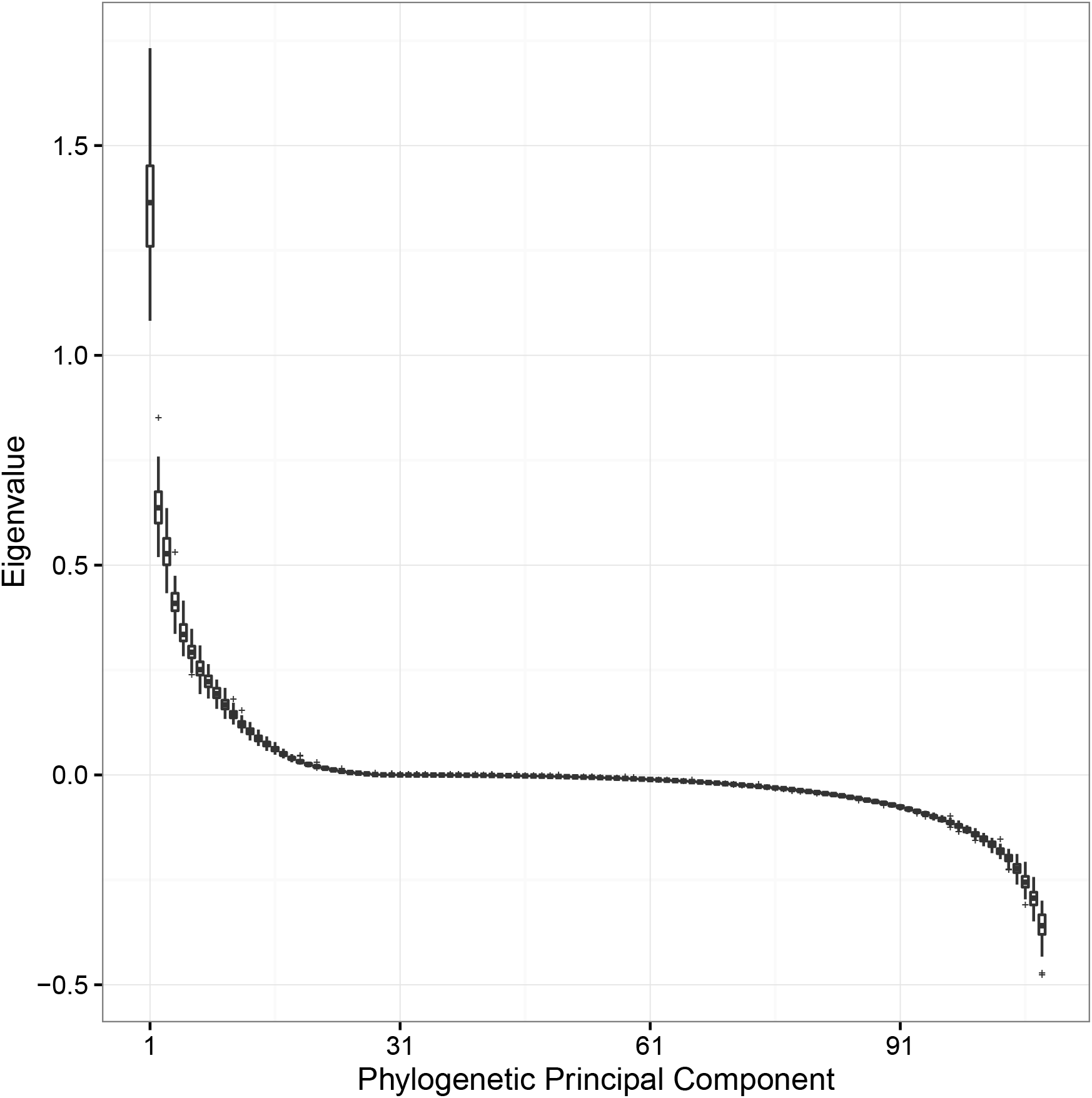
Posterior distribution of eigenvalues obtained for pPCs. Positive eigenvalues are associated with phylogenetic signal; negative eigenvalues are associated with convergence.

**Figure 4:**
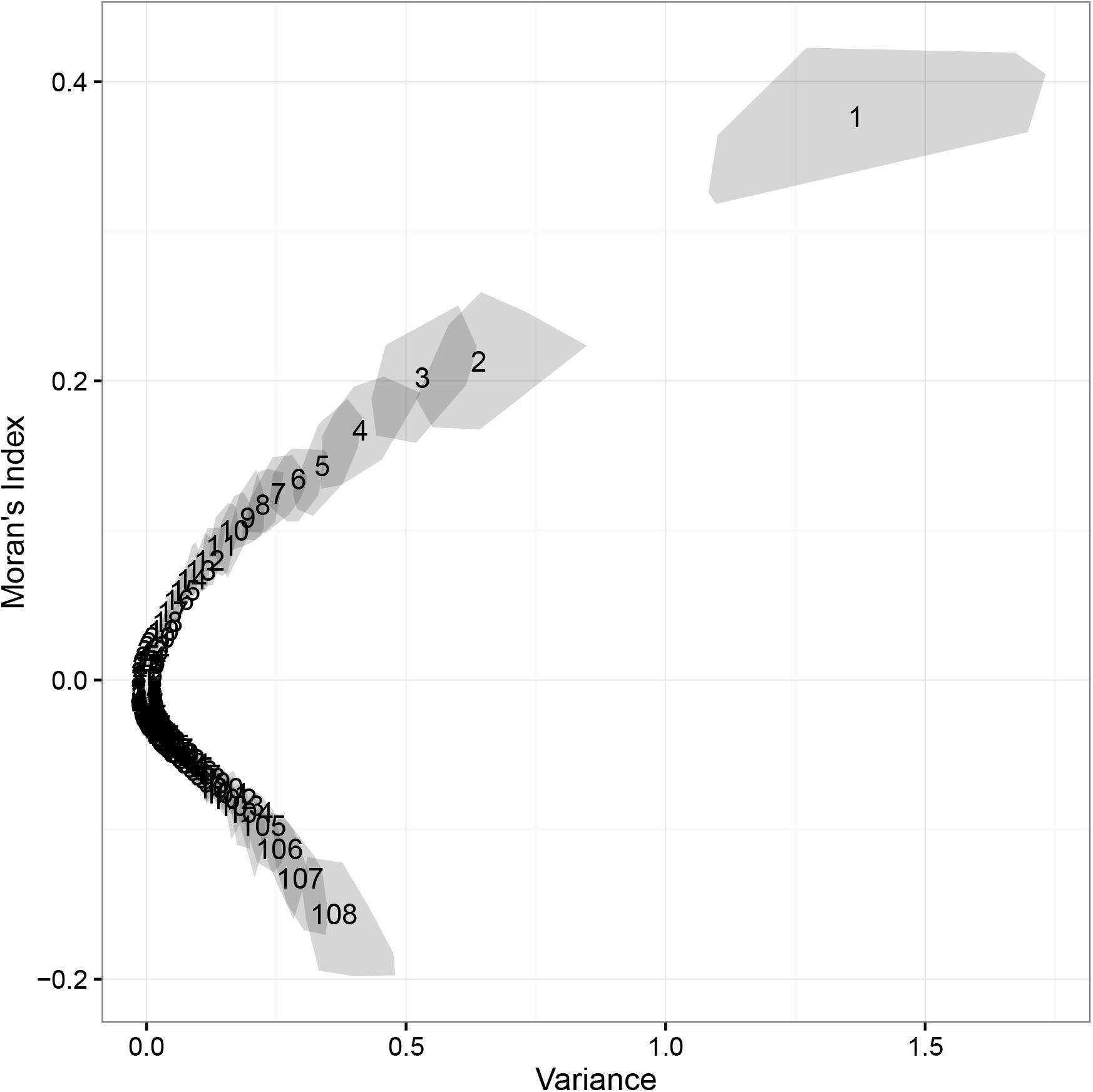
Posterior bivariate distribution of variances explained by each phylogenetic PC *versus* Moran’s Indexes estimated for each axis.

The variation in trait-specific covariance structure described by these four phylogenetic Principal Components, as captured by comparing the posterior distribution of confidence intervals for each axis using Selection Response Decomposition (Figure 5) indicates that, regardless of which axis is considered, traits with lower posterior SRD scores are usually localized in either Orbit or Basicranium, along with log Centroid Size, which represents the covariance structure associated with allometric relationships. Traits in remaining skull regions (Oral, Nasal, Zygomatic, Vault) consistently exhibit higher SRD scores, thus indicating a more stable covariance structure associated with these regions throughout Anthropoid diversification.

The overall distribution of average posterior SRD scores along the entire set of phylogenetic PCs (Figure S5) indicates that this behavior detailed on Figure 5 for G1-3 and L1 is the norm also for the remaining pPCs, that is, Orbit and Basicranial traits along with allometric relationships consistenly have lower average SRD scores than other skull regions. Notice that the explained amount of variance for intermediate pPCs (roughly from pPC 31 to 61) is very close to zero (Figure 3); thus, although matrices describing confidence intervals in these axes can be recovered, the pattern expressed by them should not be taken into account.

## Discussion

Since its conception, the hypothesis that functional interactions among morphological traits shape their phenotypic covariance structure (Olson & Miller, 1958) has been complemented with the notion that developmental interactions mediate the functional relationships among traits dynamically. This means that it is difficult to separate the relative contribution of either development or function to phenotypic integration (Cheverud, 1996a), especially considering that the structure of developmental interactions is thought to match the pattern of optimal functional interactions (Lande, 1980; Cheverud, 1984; Wagner, 1996; Jones, 2007), which further entangles both phenomena. Thus, changes in covariance structure between sister lineages should be associated with the interplay between functional and developmental interactions.

**Figure 5:**
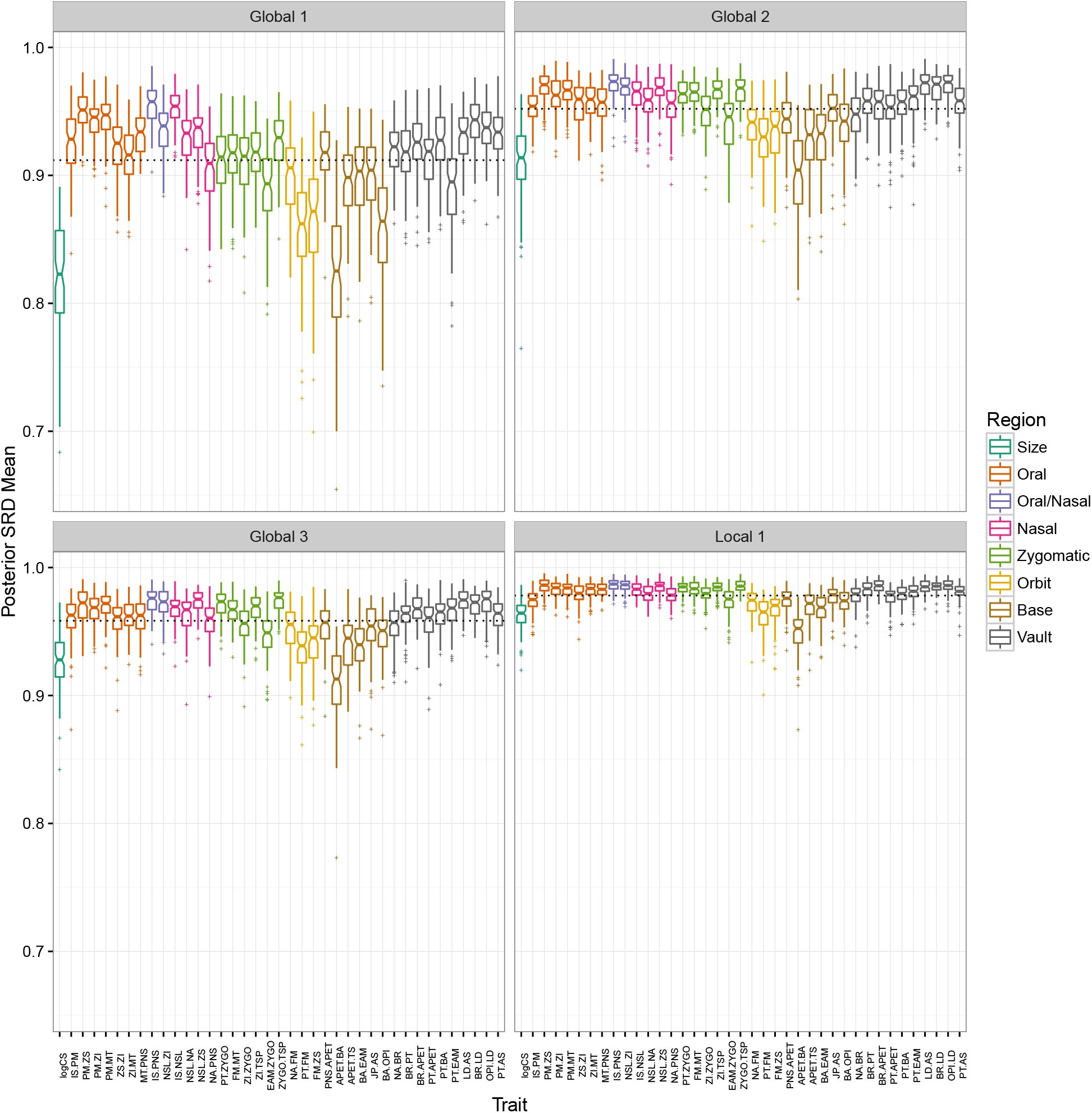
Covariance structure variation associated with the first three and last pPCs (Global 1-3 and Local 1, respectively), represented using posterior mean SRD scores. Dotted lines represent average SRD scores for each comparison. Traits are colored according to their assocation with each skull region, according to the legend.

Our results indicate that the changes in phenotypic covariance structure associated with all phylogenetic principal components follow a similar pattern with respect to changes in trait-specific covariances (Figures 5 and S5). These changes are mostly associated with either Basicranial and Orbital trait sets, along with allometric relationships, represented by the covariances between log Centroid Size and local shape variables. While the comparison between covariance matrices in the limits of each phylogenetic PC represents the overall pattern of dissimilarity in trait-specific covariance structure associated with each axis, the actual values obtained represent only a fraction of the overall divergence between Anthropoid lineages. Given any two P-matrices associated with a pair of species, the actual SRD comparison between them will be some linear combination of their divergences along each pPC axis. We now attempt to interpret these results in terms of the recent advances in developmental biology for each cranial region.

Orbital traits are mostly associated with the development of the postorbital bar in Euprimates, as opposed to Plesiadapiforms, and in Anthropoids it fully develops into the orbital cavity and postorbital wall (Ravosa & Savakova, 2004). The origins of this structure have been linked to the distribution of masticatory loadings around the comparatively large primate eye for the maintenance of a stable, forward-facing visual field even during feeding behavior (Ravosa *et al.*, 2000). Although most of the cranial Vault originates from intramembranous ossification induced by the growing brain and thus derived from the neural crest, the influence of mesoderm-derived condensations and its pattern of endochondral ossification is necessary for the proper development of the fronto-nasal and fronto-zygomatic sutures, which affect the brow ridge and both medial and lateral orbital walls (Jiang *et al.*, 2002).

In the same manner, the Basicranium originates from a set of thirteen condensations derived from both precursors, which exhibit a mosaic pattern of both endochondral and intramembranous ossification (Lieberman, 2011). Such processes occurs early during skull development, and the spatial overlap of developmental processes (Hallgrímsson *et al.*, 2009) may also explain Basicranium variation in covariance structure. Moreover, the angulation between anterior and posterior elements of the Basicranium has significantly changed during primate evolution, and such property appears to have evolved in coordination with Facial growth relative to the cranial Vault, accomodating both structures on each other (Scott, 1958; Lieberman *et al.*, 2000, 2008). Therefore, both Orbit and Basicranium are located in the boundary between domains of two precursor tissues that originate osteological elements in the skull, and their proper development is affected by both of them; furthermore, their functional significance is also associated with the accomodation of remaining structures. These characteristic properties of their development may thus be sufficient to explain their divergence among lineages in terms of covariance structure.

On the other hand, the structure of phenotypic covariances for Oral, Nasal and Vault traits remained stable through the course of Anthropoid diversification. These regions exhibit more consistent patterns of developmental processes, when compared to the Orbit or Basicranium. Both Oral and Nasal regions exhibit a great degree of interactions with soft tissue during prenatal development; the Oral region further suffers the influence of muscleskeletal interactions associated with Facial growth in postnatal development, which also contributes to its pattern of stability in covariance structure (Zelditch *et al.*, 1992; Herring, 2011; Lieberman, 2011). The cranial Vault exhibits a more regular pattern of growth, induced by the underlying brain (Lieberman, 2011; Esteve-Altava & Rasskin-Gutman, 2014). While there are prenatal developmental processes associated with the integration of both structures (Marcucio *et al.*, 2005, 2011) and postnatal muscle-bone interactions may be understood as a overall integrating factor – since the skull as whole is affected by such interactions – each of these regions is located within the bounds of precursor components, thus exhibiting a stable association between developmental processes that originate these regions and their functional aspects.

Finally, allometric relationship exhibit changes in covariance structure across phylogenetic principal components with magnitudes equivalent to Basicranial traits (Figures 5 and S5). While such relationships are explored elsewhere (Garcia *et al.*, in prep), it is noteworthy that these changes may be associated with the role of directional selection for body size, which shaped the extant diversity of New World (Marroig & Cheverud, 2005, 2010) and Old World Monkeys (Cardini *et al.*, 2007; Cardini & Elton, 2008), as well as the more complex relationships between size and shape observed within Hominidae (Ackermann & Cheverud, 2004; Mitteroecker *et al.*, 2004; Mitteroecker & Bookstein, 2008; Schroeder *et al.*, 2014). To some extent, selection for body size produces mean shape differences among lineages in a manner consistent with the ancestral allometric relationships (Lande, 1979; Schluter, 1996), but selective pressures for size may alter such relationships depending on the structure of interactions between size, shape and ontogeny (Pélabon *et al.*, 2013, 2014).

The phylogenetic distribution of changes in covariance strucuture (Figure 2) reveals an even distribution of such changes throughout Anthropoid diversification, with three different instances in which more substantial changes have occurred, in order of the estimated P-matrix diversity at each point (Figure 2a): the divergence between New World and Old World Monkeys, between Atelidae and Cebidae, and between Hominoidea and the remaining Anthropoids. Results from the phylogenetic principal component analysis (Figure 2b) recover a similar pattern, considering the contribution of these lineages to each axis of matrix variation, and the posterior distributions of the associated eigenvalues for the first, second and third phylogenetic principal components (Figure 3) indicate that these cladogenetic events can be set apart in terms of their relative contributions to P-matrix diversity. However, the apparent greater contribution of the divergence between Atelids and Cebids to P-matrix diversity may be a consequence of the phylogenetic structure imposed by either analyses, given that the actual pattern of divergence in the third phylogenetic component (Figure 2b) puts the Hominoid lineage apart not only from its sister group, Cercopithecoidea, but also from remaining lineages. Since the third pPC is not a proper constrast between sister lineages, but rather between Hominoidea and the paraphyletic grouping obtained from removing this group, the imposed phylogenetic structure of both models might reduce the contribution of this lineage to overall P-matrix diversity in either analyses.

Nonetheless, both events are overshadowed by the separation between New World and Old World Monkeys, since this divergence is associated with an unequivocally higher eigenvalue, as indicated by their posterior distribution (Figure 3). Considering that the migration from Africa to South America probably occured through the colonization of successive island environments that may have occurred in the South Atlantic Ocean during the Eocene (Oliveira *et al.*, 2009a), the effective population sizes in the ancestral population for all Platyrrhines probably plummeted during this vicariant event. Thus, the subtle differences in P-matrix diversity such separation represents may indicate that changes in the genetic architecture of cranial traits due to drift and/or founder effects (Goodnight, 2000; Brito *et al.*, 2005) may be a null hypothesis with enough explanatory power for this divergence, against an alternative hypothesis of differences in covariance structure between Platyrrhines and Catarrhines due to either directional or stabilizing selective pressures. Although these hypotheses could be tested using comparative methods, for instance (e.g. Haber, 2015), it is unclear at this point whether these methods are adequate to test hypothesis with respect to the evolution of morphological integration, given that they were conceived to model the evolution of mean phenotypes under the general assumption of constant genetic variances (Hansen & Martins, 1996). The descriptive approach we use here, focussing on comparing credible intervals between covariance matrices while incorporating phylogenetic structure has the advantage of imposing minimal assumptions. However, regardless of whether comparative methods are suited to deal with the evolution of morphological integration or not, addressing this question requires adequate sampling both within and between lineages (Melo *et al.*, 2015).

The pattern recovered by the first Local component (lower right panel of Figure 5) also describes a pattern similar to that of Global components. However, the distribution of lineages in this component scales with the logarithm of sample sizes (Figure S4), thus indicating the influence of sampling effects upon this component. Basicranial and Orbital traits exhibit overall lower covariances, both among themselves and between other regions, probably owing to the influence of multiple and disparate developmental processes; however, such lower correlation values also imply more uncertainty in their estimation. On the other hand, it is also important to consider that these directions depicted as changes in trait-specific covariance structure using the SRD method are orthogonal in the Euclidean image of the exponential-logarithm map. Thus, these components represent different aspects of covariance divergence among lineages, which in this particular case are concentrated in the same traits regardless of the phylogenetic scale considered. The association between the first Local component with sample sizes hinders its interpretation as biologically meaningful, given that Local components should be related to convergence among phylogenetically disparate lineages (Jombart *et al.*, 2010); rather, this component seems to be associated with divergence in the statistical properties of estimated covariance matrices for sister lineages.

Considering both the pattern associated with the first Local component and the prevalence of variation associated with Global components over Local components (Figures 3 and 4), the contribution of historical constraints is greater than the contribution of convergence among disparate lineages. Given that both sets of components are associated with the same structure of changes in covariance structure (Figures 5 and S5) related to some of the properties of skull development, these results highlight the overall stability of shape covariance structure in Anthropoids, in agreement with previous works using traditional morphometrics (Marroig & Cheverud, 2001; Oliveira *et al.*, 2009b).

## Conclusions

The stability of shape covariance structure in Anthropoids may be a consequence of either constraints on mammalian skull development or the prevalence of a constant pattern of functional relationships imposed by stabilizing selection; such dichotomy has already been pointed out by other authors (e.g. Marroig & Cheverud, 2001; Porto *et al.*, 2009; Oliveira *et al.*, 2009b). Here, we favor the second point of view, in light of the evidence for the evolution of genetic or phenotypic covariances under directional selection (Jones *et al.*, 2012; Melo & Marroig, 2015; Careau *et al.*, 2015) or under relaxation of stabilizing selection (Jamniczky & Hallgrímsson, 2009). Furthermore, the available comparative data (Monteiro & Nogueira, 2010; Haber, 2015) indicate that some mammal lineages, such as phylostomid bats and ruminants, have diverged in phenotypic covariance structure probably due to changes in the adaptive landscapes resulting from differences in ecological processes acting on different lineages. On the other hand, for Anthropoids, their stability in P-matrix structure may thus be possible as a consequence of stability of the functional relationships imposed over by the adaptive landscape in this lineage, as it seems that in this group, dietary shifts produced changes on body size only, at least for Platyrrhines (Marroig & Cheverud, 2010).

From an information theory point of view (Brooks *et al.*, 1989; Frank, 2009), natural selection will increase the correlation between information encoded in a population and the information that represents its environment, and the suggestion that developmental systems share properties with machine learning algorithms (Watson *et al.*, 2014) only reinforces such view. Furthermore, the regulation of developmental systems, through both genetic and epigenetic effects, may also be targeted by selection for both robustness and replicability (Hansen, 2011). These different ways of thinking about developmental and functional interactions mean that the probabilistic distribution of phenotypes, as a consequence of either genetic information or epigenetic effects, will tend to match the properties of the distribution of fitness in a particular environment. Thus, the maintenance of the same phenotypic distribution over macroevolutionary timescales indicates that the fitness distribution itself may be stable as well.

## Acknowledgements

We would like to thank G. Burin, D. A. R. Melo and M. N. Simon for comments on early versions of this manuscript. This work has been funded by CNPq (Conselho Nacional de Pesquisa e Desenvolvimento) and FAPESP (Fundação de Amparo à Pesquisa do Estado de São Paulo, grants no. 2011/14295-7 and 2011/52469-4).

